# ASD Neuro Agentic (ANA): An Agentic Framework for Feature Discovery and Cross Site Generalization in Autism fMRI Classification

**DOI:** 10.64898/2026.11.07.765076

**Authors:** Karim Rahimian, Mohammadamin Mahmanzar, Reyhaneh Bahreyni, Saleha Bayat, Lauren Higa, Kaveh Kavousi, Youping Deng

## Abstract

Classifiers of autism spectrum disorder (ASD) from resting state functional MRI rarely generalize across sites, and many reported accuracies are inflated by data leakage. We show that an agentic artificial intelligence framework, ASD Neuro Agentic (ANA), whose language model agents propose features from the literature and researcher input while fixed tools perform every calculation, can search auditably for site independent classifiers while controlling the leakage. In 1,028 participants from 17 ABIDE I centers, ANA tested 71 pipelines and selected a ComBat harmonized ensemble of tangent space connectivity models from six parcellations and 98 public connectivity derivatives. In leave one center out testing, the ensemble reached an area under the curve of 0.814 (95% CI, 0.773–0.841), outperforming ASD-DiagNet and ASD-SAENet. Predictive connectivity was mainly reduced in ASD. Agentic AI offers a systematic and efficient approach to exploring, evaluating, and identifying pipelines as model selection grows complex.

## Introduction

Autism spectrum disorder (ASD) is a neurodevelopmental condition characterized by differences in social communication and interaction, along with restricted or repetitive behaviors, interests, or activities^1,2^. ASD varies greatly among individuals and manifests in different symptoms, cognitive ability, development, and cooccurring conditions. Brain measurements also show wide variation; therefore, one biological pattern is unlikely to describe the full spectrum^1,3^. Neuroimaging studies report differences across several brain systems, including the default mode, salience, attention, sensory, frontoparietal, and social cognitive systems^4,5^. In the 2022 U.S. Autism and Developmental Disabilities Monitoring Network, ASD was identified in 32.2 per 1,000 eight year old children, or about 1 in 31, with clear differences between sites^6^. Diagnosis is still based on clinical assessment. No fMRI measure is accepted as an independent diagnostic biomarker^7^. For this reason, researchers are studying imaging features that may support, but not replace, clinical assessment^8^.

Resting state of functional magnetic resonance imaging (rs-fMRI) measures spontaneous blood oxygen level dependent (BOLD) activity without asking a participant to complete a task. This makes it useful for developmental groups with varied abilities^8^. Early computational studies of ASD mainly used functional connectivity (FC), which is usually calculated from statistical relationships between regional BOLD time series^9^. The Autism Brain Imaging Data Exchange (ABIDE) brought together 1,112 rs-fMRI datasets from 17 international sites and later expanded through ABIDE II^10,11^. This allowed multisite studies at a scale that most individual centers could not reach. At the same time, ABIDE showed a major problem. Differences in scanners, acquisition protocols, recruitment, and sites can add variation that is not related to ASD, and prediction models may learn this variation by mistake^12,13^.

Early ABIDE studies showed that whole brain FC separates ASD from controls, but with modest multisite accuracy^14^, a mixture of lower and higher connectivity^11^, and strong dependence on preprocessing, parcellation, connectivity estimation and feature selection^12^; a small set of abnormal connections did, however, generalize to independent groups^15^. Over the following decade, features moved from static FC to recurrent and deep networks^16,17^, population and interpretable graph models^18,19^, wavelet and time frequency representations^20,21,22,23^, connectome dynamics and regional genetic analyses^24,25^, and attention based and multimodal models^26,27^ (Figure S1). This greatly enlarged the space of possible pipelines while reproducibility problems became clearer: many FC differences do not replicate across sites^28,29^; small samples, class imbalance and weak external validation remain common^8,9,30,31^; and classical or simple models can match graph deep learning on ABIDE^32^. Heterogeneous functional states within ASD further suggest that no single brain signature should be expected^3^. The central question is therefore no longer whether fMRI can classify ASD, but which analysis pipeline works at a new site.

The same, BOLD data can be represented as regional activity, static or dynamic FC, time sequences, graphs, wavelet coefficients, or learned features. Each representation can also be combined with different preprocessing methods, atlases, models, and settings^9^. These choices affect each other. A method that works with one setting or random data split may fail with another model or at an unseen site^31^. Data leakage is a serious risk when feature selection, harmonization, or model tuning uses information from held out participants. A recent reanalysis of ABIDE showed that leakage can make results appear stronger than they are^33^. Participants should remain separate between training and testing. Feature selection and tuning should use training data only, and models should be tested at independent sites^12^.

Interpretation also requires caution: regions or connections that contribute to classification are not necessarily causal in ASD, and their importance may vary with preprocessing, cross-validation folds, acquisition sites and model choice ^12,13^. Likewise, the value of representations such as wavelets, which capture multiscale temporal patterns, must be established empirically against conventional connectivity under rigorous cross-site validation

Agentic artificial intelligence may help researchers examine this large search space. An agentic system can plan, use tools, evaluate results, and revise its next step instead of giving one answer. This field of practice is known as agentic science. The Virtual Lab also showed how several specialized AI agents can organize computational experiments in an iterative scientific process^34,35^.

We introduced ASD Neuro Agentic (ANA), an auditable, agentic framework for the systematic discovery of ASD classification pipelines that generalize across acquisition sites (Figure 1). Rather than relying on a single predefined model, ANA coordinates specialized agents to propose candidate methods, evaluate alternative feature representations and modeling configurations, learn from previous experimental outcomes, and refine the search based on accumulated evidence. Crucially, pipeline selection is guided by leave one center out (LOSO) validation, complemented by assessments of robustness, interpretability, and computational cost, rather than performance on random data splits. By integrating evidence guided search, iterative experimentation, and rigorous cross site evaluation, ANA provides a systematic approach to discovering and validating reliable fMRI-based ASD classifiers while addressing methodological variability, data leakage, and limited generalizability.

**Figure 1.**
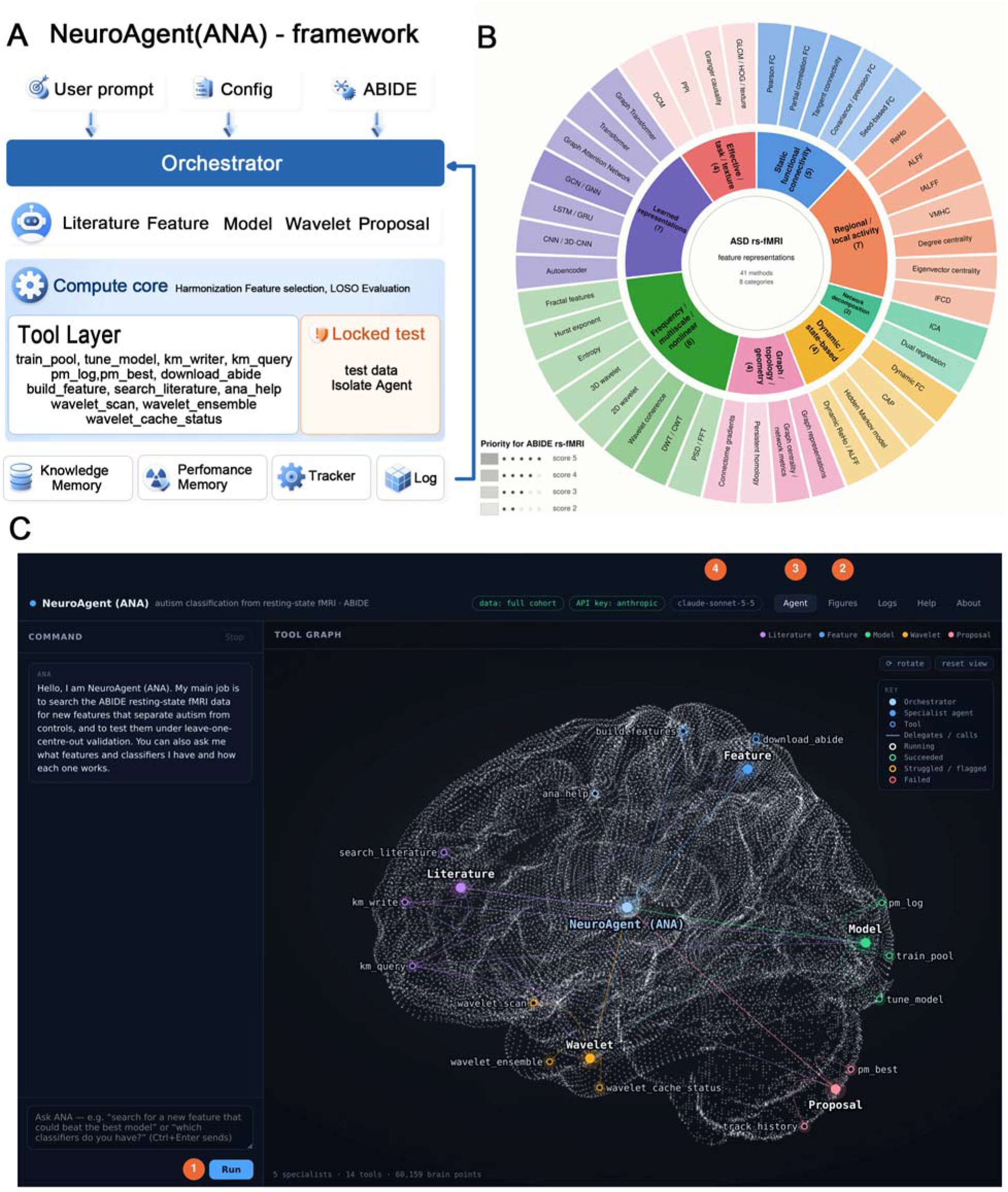
(A) Framework layers. The user prompt, configuration, and ABIDE data enter through the Orchestrator, which delegates tasks to the Literature, Feature, Model, Wavelet, and Proposal agents. In the compute core, the tool layer runs every calculation, including harmonization, feature selection, and leave-one-site-out (LOSO) evaluation. The locked test data (orange) are isolated from the agents. Knowledge Memory, Performance Memory, the tracker, and the log store the results of each run and feed them back to the Orchestrator. (B) Resting-state fMRI feature representations for ASD classification. The inner ring shows eight groups, and the outer ring shows their 41 methods: static functional connectivity (n = 5); regional and local activity (n = 7); network decomposition (n = 2); dynamic and state-based methods (n = 4); graph, topology, and geometry (n = 4); frequency, multiscale, and nonlinear methods (n = 8); learned representations (n = 7); and effective, task, and texture features (n = 4). Color marks the group, and darker shades mark higher priority scores, from 2 to 5. The score set only the order of testing and did not measure evidence strength. (C) Web interface. A request is typed in the command box and started with Run (1). The Figures tab (2) shows the study figures. The Agent tab (3) shows the live tool graph, in which the five specialist agents and 14 tools are placed on a rotatable three-dimensional brain and light up while they run. The language model serving the session is shown at (4). Example prompts and the empty trace panel are not shown.

## Results

### Autonomous search and convergence

We first examined whether ANA’s search found reproducible improvements rather than simply the best of many trials. All model selection and evaluation used leave one center out (LOSO) validation, in which every participant from the held-out center is excluded from model development.

The system tested 73 proposed pipelines. Each pipeline combined one feature representation with one classifier and received a score calculated by the tool layer under one ranking rule (Figure 2A; the best pipeline for each feature classifier pair is shown in Figure 2B and ROC curves in Figure 2C). Performance improved in only a few steps. Balanced accuracy started near 0.58, reached about 0.726 with multi atlas tangent space features and 0.741 with the final selected pipeline. This design remained the best after the system tested correlation, partial correlation, graph measures, regional activity, ICA features, volumetric features, and wavelets.

**Figure 2.**
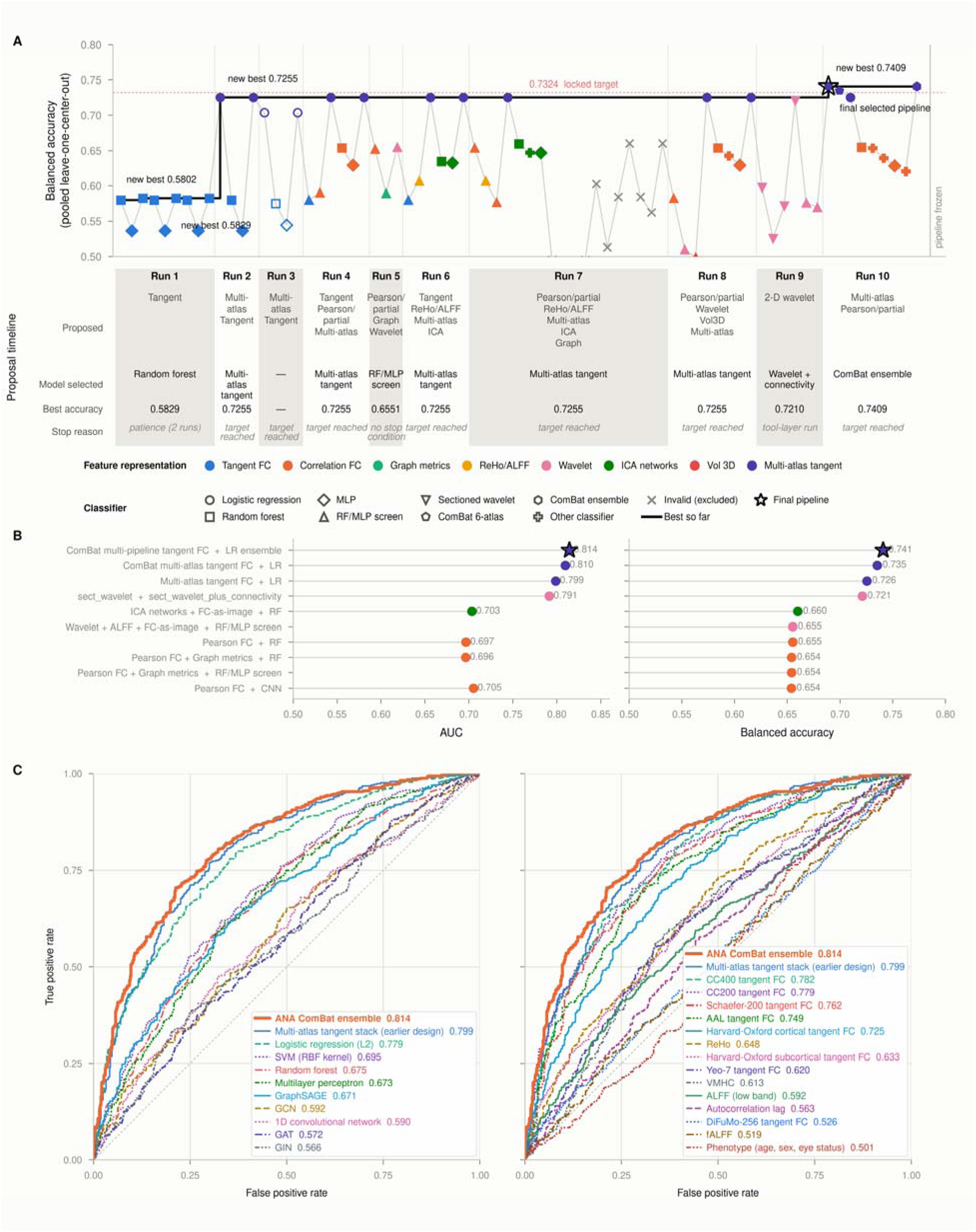
Search history of ANA model development. (A) Candidate pipelines in the order tested. The thin gray line connects tests, and the thick black step line shows the best balanced accuracy reached so far. Color shows the feature representation, and marker shape shows the model family. Filled markers are results from the full cohort (n = 1,028); open markers are from the development cohort (n = 797) and are not included in the running maximum; crosses mark evaluations excluded because of a corrupted data split. The dotted line marks the target balanced accuracy (0.7324), and the vertical line on the right marks the point at which the pipeline was frozen. The table below the plot gives, for each of Runs 1–10, the proposed representations, the model selected, the best balanced accuracy and the stopping reason (all evaluations are listed in Table S5). (B) Best result for each feature–classifier pair in the full cohort: AUC (left) and balanced accuracy (right); the star marks the final selected pipeline. (C) Pooled ROC curves of the selected ComBat ensemble compared with other classifiers (left) and with other feature representations (right); AUCs are given in the legends. Selection used pooled leave-one-center-out balanced accuracy from out-of-fold predictions. See also Table S5.

The selected ComBat ensemble drew on 104 candidate members: tangent space connectivity from six parcellations (AAL, CC200, CC400, Schaefer-200 and the Harvard Oxford cortical and subcortical atlases) and 98 ABIDE-PCP connectivity derivatives. Members were harmonized with ComBat, classified with L2-regularized logistic regression and combined by greedy forward selection on inner LOSO predictions. The contribution of each parcellation is reported in Table S2.

### Whole cohort classification performance

Across 1,028 participants from 17 centers, the selected model reached an AUC of 0.814 (Figure 3B) and a balanced accuracy of 0.74. The DeLong 95% confidence interval for AUC was 0.788– 0.840. The center level cluster bootstrap interval was 0.773–0.841 (Figure 3A). Participants from the same center are not fully independent, so the center level interval is the more cautious estimate. In 10,000 label permutations, none reached the observed performance.

**Figure 3.**
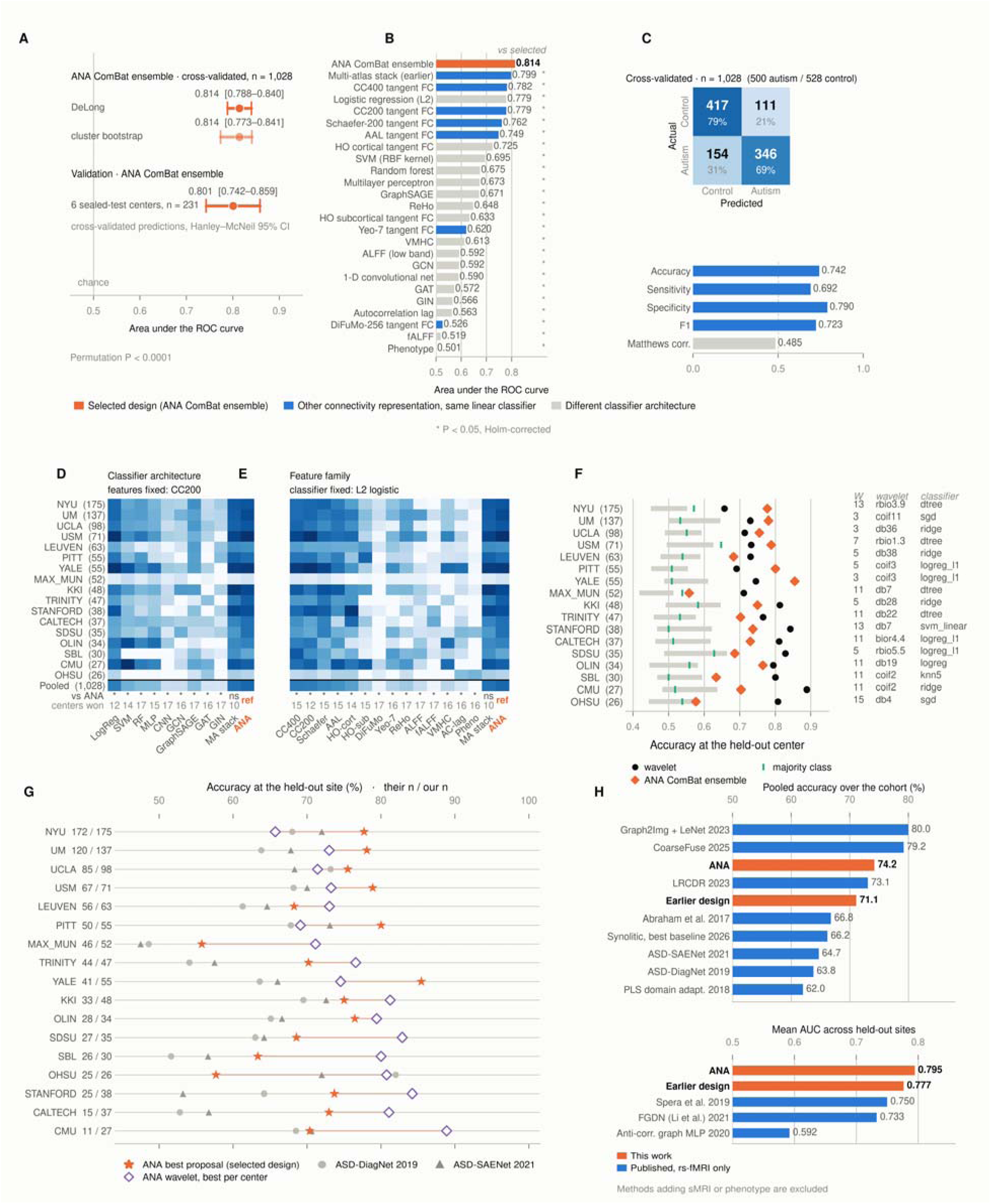
Cross-validated performance of ANA and comparison models. Results use leave-one-center-out testing across 17 ABIDE I centers (n = 1,028; 500 ASD and 528 controls). (A) The selected ComBat ensemble reached an AUC of 0.814. The DeLong interval was 0.788–0.840, the center-level bootstrap interval was 0.773–0.841, and label permutation gave P < 0.0001. On the six sealed-test centers (n = 231), its cross-validated AUC was 0.801 (Hanley–McNeil 95% CI, 0.742–0.859). (B) Ranking of 25 models by AUC. Orange marks the selected model, blue marks another connectivity representation with the same linear classifier, and gray marks other classifiers. Holm-corrected Wilcoxon tests compare each model with the selected design. (C) Confusion matrix and threshold metrics of the selected model over all 17 held-out centers (threshold 0.5). (D and E), Center AUCs after changing the classifier (D); features fixed to CC200) or the feature representation (e; classifier fixed to L2 logistic regression). (F) Wavelet results by center. Gray shows the mean ± SD. across 7,844 settings, black shows the best setting, orange shows the ComBat ensemble, and green shows the majority-class baseline. (G) Site accuracy compared with four results reported by Rahman et al. (2026). These comparisons need caution because the wavelet value is a selected maximum and the studies used different site samples. (H) Pooled accuracy over the cohort (top) and mean AUC across held-out sites (bottom) compared with published rs-fMRI-only results; orange, this work (ANA and the earlier six-atlas design); blue, published methods († as reported by Rahman et al.). Methods that added structural MRI or phenotype were excluded. See also Tables S2 and S3.

A more complex model did not give better generalization in this study. A regularized linear classifier with a suitable connectivity representation performed better than the more complex models that ANA tested. The six atlas pipeline selected in the first search phase was tested once on the six locked centers (n = 231). Its AUC was 0.768 (95% CI, 0.706–0.830), accuracy 0.701, sensitivity 0.630 and specificity 0.764. On the same 231 participants, the final ComBat ensemble had a cross validated AUC of 0.801 (95% CI, 0.742–0.859; Figure. 3C); this is not an independent test, because these centers were part of its development. Across all 1,028 participants, its accuracy was 0.742, sensitivity 0.692 and specificity 0.790 (Figure. 3C).

### Generalization across acquisition centers

Although the pooled AUC indicated overall discriminatory performance, center specific analyses revealed substantial heterogeneity, with AUC values ranging from 0.573 to 0.897 (Table 1). The highest performance was observed at USM (AUC = 0.897; 71 participants), whereas MAX_MUN showed considerably lower discrimination (AUC = 0.573; 52 participants). CMU also achieved relatively high performance (AUC = 0.824) with only 27 participants. These findings suggest that center level differences in model performance cannot be explained by sample size alone and may reflect variation in acquisition protocols, sample characteristics, or other center specific factors.

**Table 1.** Performance of the selected ComBat ensemble at each center. Out-of-fold predictions from pooled leave-one-center-out evaluation (n = 1,028). Balanced accuracy, sensitivity and specificity use a probability threshold of 0.5. Centers are ordered by AUC. See also Figure S2 and Table S1

| ***Center*** | ***n*** | ***ASD / control*** | ***AUC*** | ***Balanced accuracy*** | ***Sensitivity*** | ***Specificity*** |
|---|---|---|---|---|---|---|
| *USM* | *71* | *46 / 25* | *0.897* | *0.810* | *0.739* | *0.880* |
| *YALE* | *55* | *27 / 28* | *0.884* | *0.853* | *0.778* | *0.929* |
| *UM* | *137* | *64 / 73* | *0.859* | *0.783* | *0.812* | *0.753* |
| *KKI* | *48* | *20 / 28* | *0.859* | *0.757* | *0.800* | *0.714* |
| *STANFORD* | *38* | *19 / 19* | *0.850* | *0.737* | *0.579* | *0.895* |
| *NYU* | *175* | *75 / 100* | *0.842* | *0.763* | *0.667* | *0.860* |
| *UCLA* | *98* | *54 / 44* | *0.839* | *0.759* | *0.722* | *0.795* |
| *TRINITY* | *47* | *22 / 25* | *0.827* | *0.695* | *0.591* | *0.800* |
| *CMU* | *27* | *14 / 13* | *0.824* | *0.701* | *0.786* | *0.615* |
| *PITT* | *55* | *28 / 27* | *0.804* | *0.801* | *0.750* | *0.852* |
| *CALTECH* | *37* | *19 / 18* | *0.801* | *0.732* | *0.632* | *0.833* |
| *SDSU* | *35* | *13 / 22* | *0.801* | *0.671* | *0.615* | *0.727* |
| *OLIN* | *34* | *19 / 15* | *0.772* | *0.768* | *0.737* | *0.800* |
| *LEUVEN* | *63* | *29 / 34* | *0.749* | *0.665* | *0.448* | *0.882* |
| *SBL* | *30* | *15 / 15* | *0.711* | *0.633* | *0.667* | *0.600* |
| *OHSU* | *26* | *12 / 14* | *0.619* | *0.571* | *0.500* | *0.643* |
| *MAX_MUN* | *52* | *24 / 28* | *0.573* | *0.562* | *0.625* | *0.500* |

### Wavelet analysis

ANA also evaluated wavelet-based representations separately at each center, exploring seven section sizes (cudes with an edge of 2-15 voxel), 106 wavelet filter banks, 118 statistical descriptors, and multiple classifiers. Although some configurations achieved good performance at individual centers, no single wavelet configuration or section size consistently yielded strong performance across centers.

The center specific wavelet analysis showed encouraging classification performance for many individual datasets, although the selected configurations varied across centers (Figure 3F). For example, CALTECH used bior4.4 with L1 regularized logistic regression, whereas LEUVEN and UCLA used db38 and db36, respectively, with ridge classifiers. Other centers used different Daubechies, biorthogonal, and coiflet wavelets combined with various classifiers. AUC values ranged from 0.673 at NYU to 0.929 at OHSU, with nine of the 17 centers achieving an AUC above 0.80 and a mean classification accuracy of 76.8% across centers. These results suggest that wavelet based features can provide useful discriminatory information for individual datasets (Table S3).

These center specific results do not show that the selected configurations would generalize to unseen centers. The connectivity model performed better than the mean wavelet configuration at most centers (smallest difference at CMU), and even wavelet configurations selected with held out labels did not surpass it on the full cohort. Wavelet features thus supported classification within individual datasets, whereas connectivity features gave stronger performance overall.

### Motion and scanner effects

Head motion differed between the two groups and had a weak link with diagnosis (Figure 4A– D). Mean framewise displacement (FD), a measure of how much a participant’s head moved between consecutive fMRI volumes, was higher in the ASD group than in controls (0.156 versus 0.103 mm; Cohen’s d = 0.36; Mann–Whitney P = 1.1 × 10−9). Model predictions were also related to motion in the full sample (Spearman ρ = 0.43, P = 5.5 × 10−47) and within centers (median ρ = 0.31). However, motion did not explain the imaging result. After matching ASD and control participants on motion within each center (0.2-SD caliper, i.e. a pair was accepted only if mean FD differed by at most 0.2 SD of the cohort, about 0.03 mm; n = 732; group difference P = 0.82), motion alone fell to an AUC of 0.503, while the connectivity model kept an AUC of 0.794 (95% CI 0.745–0.832). A model with age, sex, and motion reached an AUC of 0.599. Adding the imaging decision value (the imaging model’s out of fold prediction expressed as log odds) increased it to 0.813 (Figure 4B). Age and sex alone had little predictive value. Scanner effects were much stronger: Schaefer-200 connectivity predicted acquisition center with a balanced accuracy of 0.588, compared with a 17 class chance rate of 0.059. Motion contributed to the prediction but did not account for it. The strong scanner signal supports the use of LOSO evaluation.

**Figure 4.**
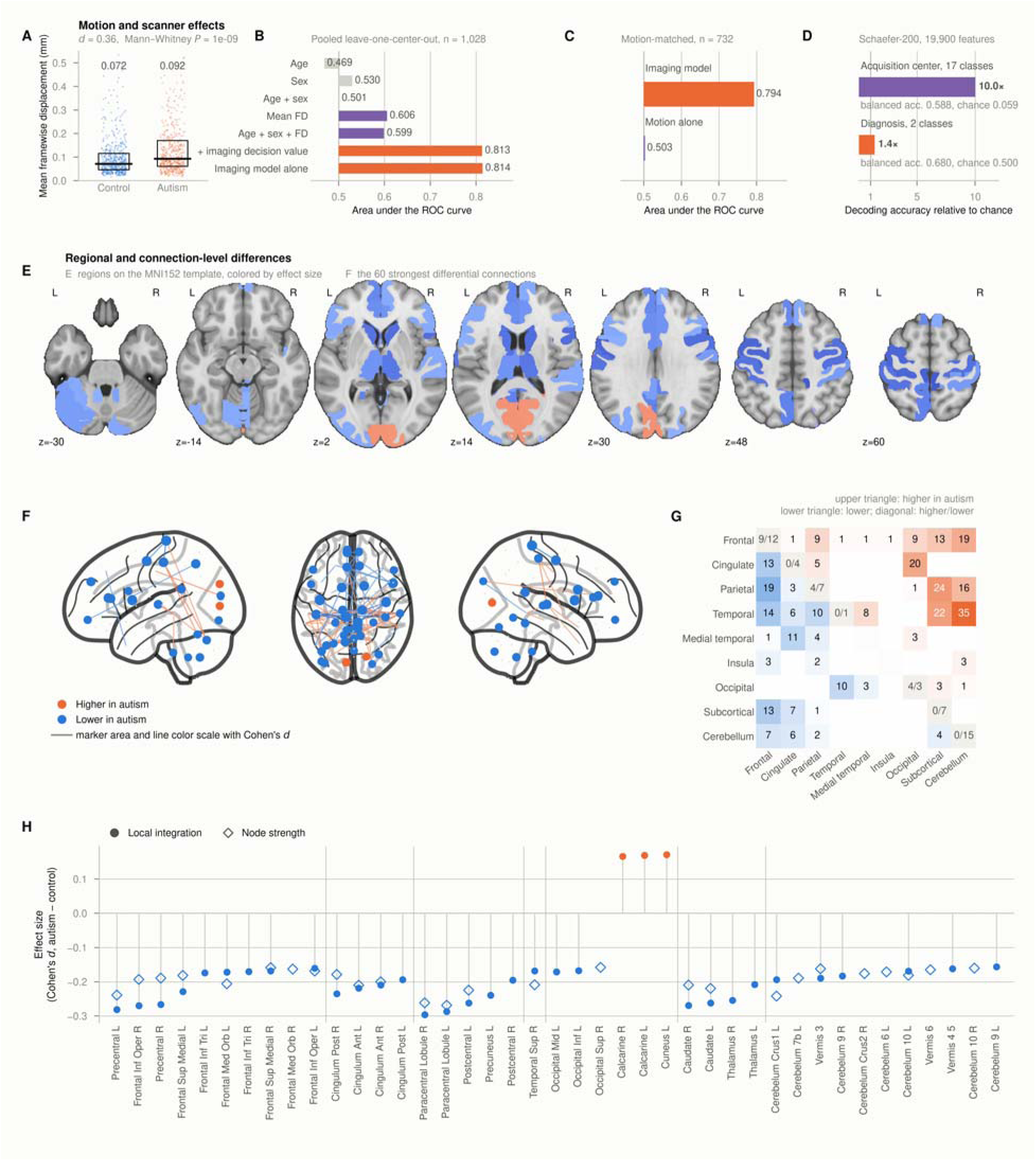
Motion and scanner effects do not fully explain classification performance, and autism-associated connectivity differences are predominantly reductions. ABIDE I included 1,028 participants (500 with autism and 528 controls) from 17 centers. Unless otherwise specified, predictions were evaluated using leave one center out cross validation. (A) Mean framewise displacement (FD) was higher in the autism group than in controls (0.156 vs. 0.103 mm; Cohen’s d = 0.36; P = 1.1 × 10). Boxes represent medians and interquartile ranges. (B) Imaging achieved an AUC of 0.814, compared with 0.599 for the combined age, sex, and FD-based model. Adding the imaging decision value yielded an AUC of 0.813. (C) Within-center motion matching retained 366 participant pairs with balanced FD (P = 0.82). Motion-only classification yielded an AUC of 0.503, whereas imaging retained an AUC of 0.794 (95% CI, 0.745–0.832), despite a correlation between imaging predictions and FD (Spearman ρ = 0.43). (D) Connectivity features decoded center identity more strongly than diagnosis relative to chance (10.0-fold vs. 1.4-fold; five-fold cross-validation), highlighting the importance of evaluation across held-out centers. (E–H) Group differences were adjusted for age, sex, FD, and center, with Benjamini–Hochberg correction applied separately across 116 AAL regions and 6,670 connections; 735 connections were significant. (E) Significant regional differences mapped onto the MNI152 template. (F) The 60 strongest differential connections, with node area proportional to |Cohen’s d|. (G) Lobe-to-lobe connection counts, with increases above and reductions below the diagonal. (H) Effect sizes for 41 significant regions. Orange indicates higher and blue lower values in the autism group. Mean-normalized ALFF showed no significant regional differences (minimum q = 0.83). The three small connectivity increases in early visual cortex may reflect the effects of prior global signal regression.

### Regional and connection level differences

We studied regional activity and connectivity while controlling for age, sex, mean FD, and acquisition center. Normalized mean of ALFF did not differ between ASD and control participants in any of the 116 AAL regions after correction (smallest q = 0.83; largest |d| = 0.14). Connectivity measures showed clearer differences. Local integration differed in 34 regions, and 31 were lower in ASD. Node strength was lower in all 26 significant regions (Figures 4E and 4h). The largest regional effects involved the bilateral paracentral lobules, precentral gyri, right inferior frontal operculum, right caudate, and left cerebellar Crus I, but all effects were modest (|d| ≤ 0.30). Three regions in early visual cortex had higher local integration. At the connection level, 735 of 6,670 connections remained significant after correction (Figures 4E and 4G). Both higher and lower connectivity were present. The main result was that group differences were clearer in relationships between regions than in the amplitude of local activity.

Regional connectivity strength was mainly lower in ASD (Figure 5A and 5B). Across the six parcellations, 173 regions passed correction: 25 of 116 in AAL-116, 41 of 200 in CC200, 61 of 392 in CC400, 38 of 200 in Schaefer-200, one of 48 in Harvard–Oxford cortical, and seven of 21 in Harvard–Oxford subcortical. Of these regions, 170 had lower strength in ASD. After mapping the findings to common anatomical names, 56 areas appeared in at least one atlas, 30 in at least two, 19 in at least three, and 11 in four or more. The left medial superior frontal gyrus appeared in five of the six atlases (d = −0.19, q = 0.017). Ten other areas appeared in four atlases (Figure 5C). The effects were small, and no shared area had |d| above about 0.30.

**Figure 5.**
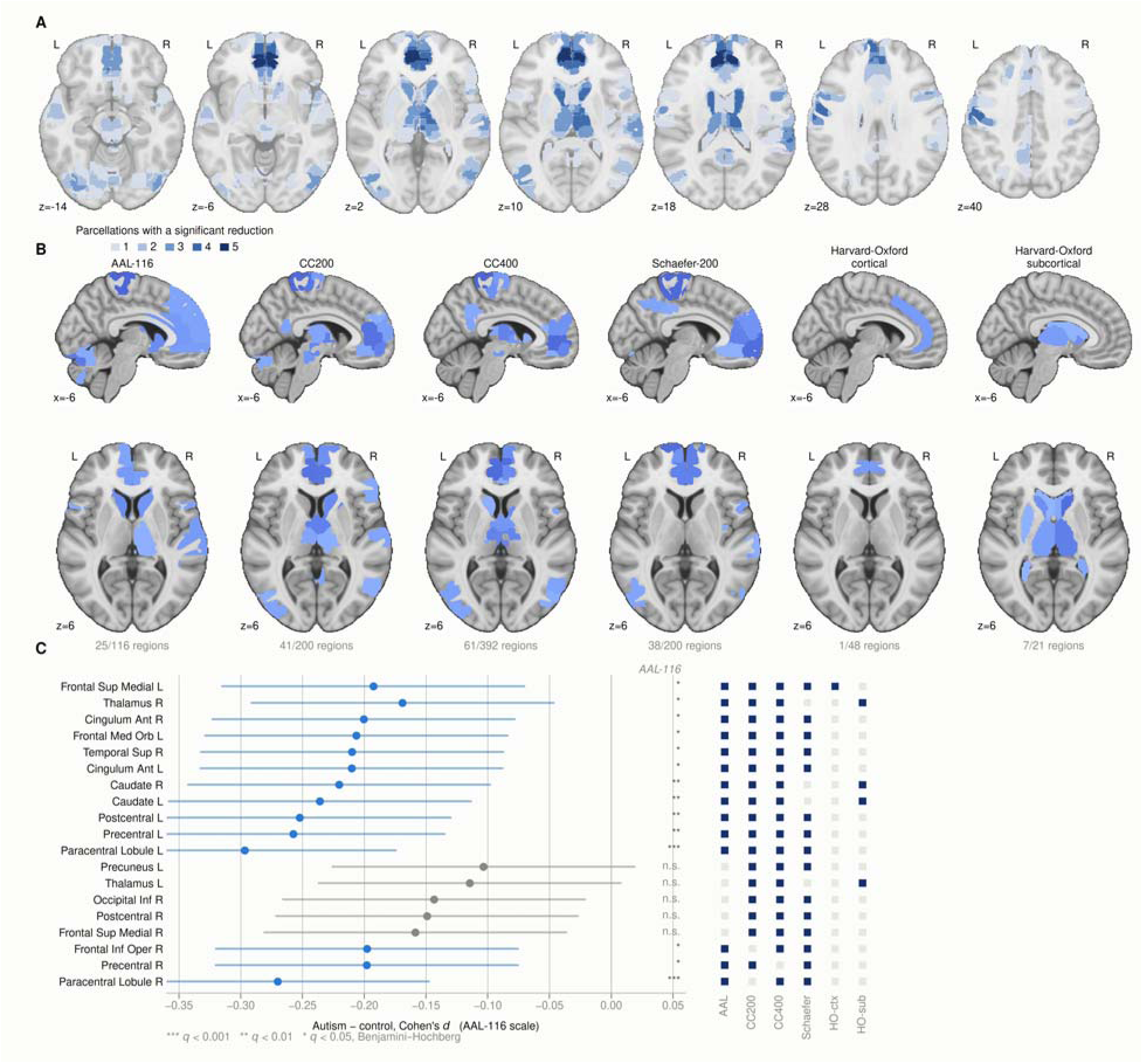
Regional connectivity strength in ASD and control participants. Connectivity strength is the mean absolute Fisher-z correlation from one region to all other regions in the same atlas (n = 1,028; 500 ASD and 528 controls; 17 centers). Each regional model included diagnosis, age, sex, mean framewise displacement, and center. Benjamini–Hochberg correction was applied within each parcellation. (A) Agreement map on the MNI152 template. Each voxel shows how many parcellations place it in a region with lower connectivity strength in ASD. (B) Results for each parcellation. Blue shows lower strength in ASD and red shows higher strength. Of 173 significant regions, 170 were lower in ASD. (C) Nineteen anatomical areas where at least three parcellations agreed on a reduction. Points show AAL-116 values, and bars show 95% intervals. The grid shows which parcellations supported each area. The effects range from d = −0.10 to −0.30. Both lateral ventricles were also identified by Harvard–Oxford subcortical; these are non-neural findings and probably reflect remaining nuisance variation.

Atlas choice affected prediction more than it affected the anatomical pattern. Regional effect maps from the four whole brain parcellations were strongly correlated (Spearman ρ = 0.73–0.87; mean 0.82). No atlas produced clearly larger effects or a higher share of significant regions. With ComBat harmonization, prediction AUC was 0.796 for CC400, 0.795 for CC200, 0.740 for Harvard Oxford cortical and 0.636 for Harvard Oxford subcortical (Table S2). Averaging the six harmonized atlases gave 0.810, and the final ComBat ensemble reached 0.814. Finer atlases may add prediction information without changing the general anatomical pattern.

Three points limit the interpretation. First, the CPAC filt_global pipeline includes global signal regression, so lower connectivity is relative to this preprocessing choice. Second, the Harvard– Oxford subcortical atlas identified both lateral ventricles. Ventricular signals are not neural signals, so these results probably reflect remaining physiological or scanner variation. Third, the regional effects were small. Similar patterns across atlases support spatial consistency, but the large overlap between groups means that no single region should be treated as a diagnostic marker.

## Discussion

This study shows that an autonomous, leakage controlled search can identify an accurate and interpretable approach to autism classification from resting state fMRI. Across 1,028 participants in ABIDE I^36^, the selected model reached an AUC of 0.814 (95% CI 0.773–0.841) across 17 acquisition centers (Figure 3A and 3B). The mean held out site AUC was 0.795 for the full cohort, and the earlier six atlas design reached 0.777 on the quality controlled subset (Figure 3H), higher than the fMRI only LOSO results identified in earlier studies^37–39^. With the earlier design, on the same participants and tangent space features used by Abraham et al.^12^, accuracy improved from 66.8% to 71.1%, a gain of 4.3 percentage points (Figure 3H). Full cohort accuracy (74.2%) also exceeded several published approaches^40–42^. Our proposed model achieved 74.2%, outperforming LRCDR (73.1%). However, CoarseFuse^43^ (79.2%) and Graph2Img^44^ (80.0%) reported higher values, although their evaluation settings and metrics differ from those used in our study. Together, these comparisons support the strength of the model under site independent evaluation.

An important strength is the validation beyond model development. Six centers, comprising 231 participants, were kept sealed until the first phase pipeline was fixed, and its single evaluation gave an AUC of 0.768. The result also remained strong after accounting for head motion, a known influence on functional connectivity^45^. Participants with autism had greater mean FD than controls (0.156 versus 0.103 mm), and motion alone predicted diagnosis with an AUC of 0.606. After matching participants by motion within centers, the imaging model retained an AUC of 0.794, while motion alone approached chance (Figure 4C). Adding the imaging score to age, sex and motion raised AUC from 0.599 to 0.813 (Figure 4B). Thus, the model captures connectivity information beyond these measured factors, while leaving room to examine other possible confounds.

Results across centers offer another useful perspective. The selected model, applied unchanged at every center without center specific selection, averaged 72.2% accuracy and outperformed ASD-DiagNet (63.8%) at 16 of 17 centers (P = 0.003) and ASD-SAENet (64.7%) at 15 of 17 (P = 0.002; Figure 3G). Scanner effects nevertheless remained strong: acquisition center could be identified at about ten times chance performance, compared with 1.4 times for diagnosis (Figure 4D). This difference helps explain variation between sites and supports holding out entire centers instead of using random participant splits^14,46^. The center specific results illustrate local potential, whereas the unchanged model provides the stronger test of transfer across scanners.

The sectioned wavelet analysis reached a higher mean site level accuracy (76.8%) when the best configuration was selected separately for each center, and it outperformed ASD-DiagNet at 14 centers and ASD-SAENet at 15 (both P < 0.001). The two results answer different questions, however. The connectivity model was fixed in advance and shows how one unchanged classifier performs at a new scanner, whereas each wavelet result is the best of 7,844 configurations. Pooled over all participants, the best wavelet model reached an AUC of only 0.643, compared with 0.814 for the connectivity model; wavelet accuracy rose as center size fell (Spearman ρ = −0.78) and, at 14 of 17 centers, did not exceed the maximum expected from model selection alone (Figure 3F). The fairest comparison with previous work therefore uses the fixed model, which improved mean accuracy by 7.5–8.4 percentage points over both published methods.

The model search also showed that greater complexity did not improve prediction. GraphSAGE, GCN, GAT, GIN, a convolutional neural network and a multilayer perceptron had AUCs 0.14– 0.25 lower than the selected model on identical cross validation folds (Holm-adjusted P = 0.012; Figure 3B and 3D). In the earlier six atlas design, evaluating 16,278 combinations across 14 feature families improved AUC by only 0.0006 (P = 0.79), and the finest atlas, DiFuMo-256, performed close to chance (Figure 3B and 3E). These findings agree with studies favoring regularized models for functional connectivity ^47,48^. Complex architectures have larger hyperparameter spaces, and our search budget may not have been equally sufficient for each, so the results show that they did not outperform the linear model under these settings, not that they are inherently unsuitable. Short scans and many connections relative to sample size may also limit the benefit of flexible models.

The ComBat ensemble integrates a simple linear classifier with complementary representations of brain connectivity derived from multiple parcellations and preprocessing pipelines. Tangent-space features capture covariance patterns within a shared representation space ^49^, while incorporating AAL^50^, CC200 and CC400^51^, Schaefer-200 ^51^, and two Harvard–Oxford atlases reduces dependence on any single brain parcellation. This integration improved AUC by 0.032 compared with the best-performing single atlas without harmonization (CC400; AUC = 0.782; 95% CI for the difference, 0.013–0.044; Holm-adjusted P = 0.012). It also yielded an AUC improvement of 0.016 over the six-atlas average, although the corresponding 95% CI included zero (−0.001 to 0.028; one-sided P = 0.032; Figure 3B and 3E). Although predictive performance varied across atlases, their regional patterns showed substantial agreement (Spearman ρ = 0.73– 0.87; Figure 5). These findings suggest that combining multiple atlases may capture complementary aspects of shared connectivity signals that are represented differently across parcellations. Furthermore, because each atlas contributes an independent representation, additional parcellations can be evaluated against the existing ensemble without repeating the entire model-search procedure.

The ANA adds value beyond the final classifier, extending recent work on autonomous scientific experimentation^52^. Its Knowledge Memory and Performance Memory store hypotheses, tested models and their results, so later experiments can build on earlier findings. All metrics are calculated by deterministic code, held out labels remain inaccessible to the agents, and proposals and tool calls are logged. Independent runs reproduced the selected result to four decimal places. This separation between automated exploration and fixed evaluation makes the search easier to audit and repeat, an important quality for multi site neuroimaging research.

The regional findings further support the relevance of the imaging signal. After adjustment for age, sex, motion and center, 170 of 173 significant regions had lower connectivity strength in autism (Figure 5A and 5B), and the left medial superior frontal gyrus appeared in five of six atlases (Figure 5C). Of the 6,670 AAL connections, 735 differed between groups after correction (Figure 4E-4H). Reduced coupling in autism was concentrated between frontal and parietal cortex, within the cerebellum and between frontal and temporal cortex, consistent with long range fronto posterior under connectivity, whereas increased coupling clustered in temporal cerebellar and subcortical pairs. The strongest single effects involved thalamic connections, in line with altered thalamocortical connectivity. These patterns agree with reports involving default mode, frontotemporal, sensorimotor, thalamocortical and striatal systems^53–57^.

Some aspects merit further study. Analyses used ABIDE I only, and the regional analyses used one preprocessing pipeline, so testing on ABIDE II will be important for external validation^58^. The best center specific settings were selected from several evaluated candidates and should be seen as exploratory estimates, not guaranteed accuracy at new sites. ComBat harmonization also used unlabeled scans from each held out center, so the model needs a batch of scans from a new center before it can be applied. Finally, global signal regression was applied during preprocessing, so connectivity findings are relative to the removed global signal. These considerations provide clear directions for future work; in particular, an independent test of the final model on new centers is still needed.

Overall, ANA, which separates feature proposals from fixed numerical computation, identified a ComBat harmonized ensemble of tangent space connectivity models with an AUC of about 0.81 across 17 centers under LOSO evaluation, outperforming the alternatives evaluated here. Functional connectivity predicted better than regional amplitude and wavelet features, and harmonization and parcellation ensembles mattered more than classifier complexity. The contribution is twofold: better prediction within the evaluated search space, and a reproducible, leakage-controlled framework that links every proposal, successful or not, to its evaluation. It provides a basis for further feature discovery and independent validation on datasets such as ABIDE II.

## Methods

### Participants and data

We obtained resting state fMRI and phenotypic data from ABIDE I (RRID:SCR_003612; https://fcon_1000.projects.nitrc.org/indi/abide). The release contains 1,112 datasets (Table S1) from 539 participants with ASD and 573 typically developing controls across 17 acquisition centers^36^. We used C-PAC (RRID:SCR_000862) preprocessed data from the filt_global pipeline^59^. This pipeline includes motion correction, nuisance regression, temporal filtering from 0.01 to 0.10 Hz, and global signal regression^60,61^. The classification model also used ABIDE-PCP connectivity derivatives from four preprocessing pipelines (C-PAC, CCS, DPARSF and NIAK), with and without band pass filtering and global signal regression (NIAK filtered only), in seven atlases (AAL, CC200, CC400, Dosenbach-160, EZ, Harvard–Oxford and Talairach–Tournoux). The regional and connection level analyses used the C-PAC filt_global data only. Diagnosis was defined by DX_GROUP, with ASD as the positive class. We excluded 77 participants without the required data, two with missing or unreadable archives, and five who were not in the feature cache. The final sample included 1,028 participants: 500 with ASD and 528 controls (Figure S2). No a priori power analysis was performed; the sample comprised all eligible ABIDE I participants. Sex was included as a covariate in ComBat harmonization and in all regional analyses, but analyses were not stratified by sex because 84.8% of participants were male. ABIDE I is a deidentified public dataset: each contributing site obtained approval from its local institutional review board and written informed consent (or assent with parental consent) from participants, and no additional ethical approval was required for this secondary analysis.

### Site grouping and data partitions

We grouped participants by acquisition center instead of the original ABIDE site labels. Leuven, UCLA, and the University of Michigan each had two samples collected with the same scanner. We combined these samples so that the same scanner could not appear in both training and test data. This reduced 20 site labels to 17 centers (Figure S2B). Before model development, we reserved six centers with 231 participants as a locked test set. This group included 108 participants with ASD and 123 controls. The other 11 centers, with 797 participants, formed the development set for the first search phase. The pipeline selected in that phase was scored once on the locked set. The search then continued on all 1,028 participants with LOSO validation, which produced the final model; the locked set is therefore not an independent test of the final model. This center disjoint design tests the model on unseen acquisition centers^12,62^.

### Agentic framework

The ANA framework has five operational layers: input, agents, tools, a deterministic compute core, and persistent state (Figure 1A). The input layer defines the research question, user queries, keywords, ABIDE I data, random seed, center grouping, evaluation rules, and stopping criteria. The agent layer has five parts: the Orchestrator, Literature Agent, Feature Agent, Model Agent, Wavelet Agent and Proposal Agent. Each agent can use only an approved set of tools and must return structured output. The tool layer runs fixed Python functions for literature retrieval, data access, feature building, wavelet analysis, model training, optimization and memory access. Language models can guide the search, but they do not calculate numerical results. At each round, the Orchestrator reviews the study goal, literature findings, tested settings, and stored performance. It then chooses the next analysis task.

The Literature Agent finds feature ideas supported by published work. The Feature Agent turns these ideas into feature settings that the software can run. The Model Agent tests classifiers and adjusts their parameters when needed. The Wavelet Agent searches wavelet-based representations built directly from the preprocessed 4D fMRI images. It combines, the section size (cube with an edge of 3-15 voxels; seven sizes) with the wavelet filter (106 options), computes a fixed set of 118 features for each section, tests several classifiers on each combination, and reports performance both pooled and for each center. The Proposal Agent writes the end-of-run report from the experiment history and Performance Memory; the system overwrites the report’s numeric fields with the tracked values. After each round, the system stores the literature source, feature settings, model settings, performance, and provenance record. Earlier results then guide later experiments. The search stops when it reaches the target, shows too little improvement, meets a stopping rule, or uses the available computing budget. The Orchestrator does not calculate any score; it reads numerical results from the compute core and Performance Memory (PM). As Reporter, the Orchestrator also writes the run report from the experiment history and Performance Memory; the system replaces every number in that report with the tracked value

### Literature search and feature proposals

The Literature Agent searched a wider set of representations than could be tested in one run. It turned published methods into source based feature ideas for the Feature Agent. It did not assume that functional connectivity, wavelets, or another representation would be best. Search queries combined ASD and ABIDE terms with rs-fMRI and a representation term. A typical form was: (autism OR ASD OR ABIDE) AND (resting state fMRI OR rs-fMRI) AND “representation term”.

We organized the search terms into eight groups (Figure 1B): static functional connectivity; regional and local activity; network decomposition; dynamic and state based measures; graph, topological, and geometric methods; frequency, wavelet, and nonlinear measures; learned representations; and a context group for effective connectivity, task interactions, and general image descriptors. The list included common rs-fMRI measures such as Pearson correlation, ReHo, and fALFF, as well as tangent space connectivity, dynamic FC, hidden Markov models, graph measures, wavelets, autoencoders, CNNs, recurrent networks, GNNs, and transformers. Table S4 lists the 41 methods with their groups and priority scores.

Each literature based idea needed at least one source that could be checked. We compared each claim with the abstract linked to its identifier before adding it to the provenance record. A priority score only decided the order of testing. It was not a measure of evidence strength and did not predict model performance. A score of 5 described a common rs-fMRI method with direct relevance to ABIDE. Lower scores described methods that were more exploratory or needed more caution.

### Feature computation

The Feature Agent chose representations from the literature based ideas, but fixed code calculated all feature values. This prevented the language model from making calculations or creating unsupported features. Features were calculated separately for each participant. Any value that depended on the full sample was fitted inside the cross validation process with training participants only. Connectivity features included Pearson correlation as a full connectome baseline^63^, partial correlation for conditional regional relationships^48,64^, and tangent connectivity, which maps covariance matrices into a Euclidean tangent^47–49^. The tangent space reference was fitted in each training fold and then applied to held out participants to avoid leakage^65^. Regional features included ReHo for local BOLD synchrony^66^ and ALFF/fALFF for low frequency BOLD amplitude^67^.

Other feature groups described network structure and time frequency information. We used graph measures such as node degree, clustering, and network efficiency, together with graph inputs for graph learning models^18,68,69^. Wavelet and frequency features described non stationary BOLD activity at several time scales. They included discrete and continuous wavelets, wavelet coherence, 2D and 3D wavelet features, and Fourier power measures^20,70,71^. The wider search also covered ICA, dual regression, dynamic FC, co activation patterns, hidden Markov models, connectome gradients, entropy, fractal measures, and learned features^72–76^. We tested a method only when a fixed implementation was available. Other methods remained ideas for future work. Features were mainly standardized within each center. ComBat harmonization used acquisition site as the batch and sex, age and mean FD as covariates, never diagnosis. In each outer fold it was fitted on the training centers together with the unlabeled scans of the held out center; the inner folds used for member selection saw training centers only^77,78^.

### Wavelet features

We incorporated wavelet based representations to investigate multiscale patterns in fMRI signals. Rather than relying on a single predefined feature representation, ANA autonomously constructed and evaluated candidate wavelet feature representations by coordinating spatial sectioning, wavelet filter bank selection, and statistical feature extraction. For each preprocessed 4D fMRI image, voxel space was cut into a fixed grid of W × W × W cubes (W = 3, 5, 7, 9, 11, 13, or 15 voxels), and the voxel by time matrix of each section received a single level two dimensional wavelet transform. At each segmentation setting, it evaluated 106 wavelet filter banks drawn from several wavelet families. For each resulting representation, 118 statistical descriptors were calculated from the wavelet coefficients, including minimum, maximum, mean, standard deviation, variance, entropy, and other distributional measures. These components were assembled into candidate feature representations for downstream classification. The wavelet search combined three choices: the section size, the wavelet filter, and the classifier. Several classifiers were then tested. The system did not begin with one preferred wavelet or section size. For every filter bank, it found the best combination of section size, features, and classifier. This allowed us to study the wavelet basis while considering its interaction with section size and the classifier.

### Model selection and validation

The Model Agent tested classifiers in two settings. In the per site setting, performance was measured separately at each acquisition center. This showed whether a method behaved in a similar way across scanners, protocols, and participant groups. In LOSO testing, one center was removed from model development, the model was trained on the other centers, and the removed center was used only for evaluation. This was repeated for every center. All learned steps, including scaling, harmonization, tangent space fitting, feature selection, and parameter tuning, used training data only. The agent chose which settings to test, while the deterministic compute core fitted models and calculated metrics. Performance Memory stored the results for later decisions and final model selection.

After the first search phase, ANA fixed the pipeline selected at that point (the six atlas tangent space average), fitted it on the full development set, and applied it once to the locked test set. The agents could not access the locked test data or labels. Every test attempt was recorded with the model settings, time and code fingerprint. The main outcome was area under the receiver operating characteristic (ROC) curve, with balanced accuracy as a second main outcome. We also report sensitivity, specificity, positive and negative predictive value, F1 score, and Matthews correlation coefficient at the chosen threshold. Raw accuracy was not a main measure because class balance differs between ABIDE centers. No randomization was involved because this is an observational analysis; cross validation folds were defined by acquisition center, and a fixed random seed (42) was used for all stochastic steps. Investigators were not blinded to diagnosis, but the agents had no access to held out labels. The study is reported following the TRIPOD statement.

### Tool layer and memory

A deterministic tool layer carried out all computations through an in process Model Context Protocol (MCP) server. Fourtheen typed tools supported literature retrieval, data access, feature construction, wavelet analysis, model training, optimization, and memory access (Figure 1A). The web interface adds a fourteenth tool, ana_help, which answers questions about ANA. Each agent could call only its approved tools: the Literature Agent retrieved papers and wrote to Knowledge Memory; the Feature Agent obtained data and built features; the Model Agent trained and tuned classifiers and logged results to Performance Memory; the Wavelet Agent scored wavelet representations and ensembles; and the Proposal Agent read the best recorded run and the run history. Agents selected tools but did not calculate metrics. Locked split creation and locked test evaluation were not available in the agent tool list, so agents could not access held out labels. The system recorded every tool call for audit purposes.

The system used separate stores for knowledge, performance, provenance, and reusable files. Knowledge Memory (KM) held literature based ideas and earlier findings. Performance Memory (PM) held model settings and the official scores used for selection and stopping. Other stores kept the experiment history, provenance logs, cached features, and the fixed locked split. This design allowed the system to reuse earlier results while keeping a record of each decision.

### Software and use of large language models

We built the framework in Python 3.11.15 (RRID:SCR_008394) with NumPy 2.4.6 (RRID:SCR_008633), SciPy 1.17.1 (RRID:SCR_008058), scikit-learn 1.9.0 (RRID:SCR_002577), Nilearn 0.13.1 (RRID:SCR_001362), NiBabel 5.4.2 (RRID:SCR_002498), pandas 3.0.3 (RRID:SCR_018214), PyWavelets 1.8.0 (RRID:SCR_024421) and Matplotlib 3.11.0 (RRID:SCR_008624). ChromaDB 1.5.9 stored Knowledge Memory, and SQLite stored Performance Memory. Language model requests were sent through the Anthropic Messages API. No model weights were stored or changed locally. Claude Opus handled orchestration, Claude Sonnet handled specialist reasoning and Claude Haiku handled initial literature screening. The agent loop ran in the application code rather than in a separate agent platform. Each request included the agent instructions, task, and only the tool schemas allowed for that agent. When a model requested a tool, a local Python function ran and returned the result. This allowlist limited tool access for each agent. Temperature was set to zero, and each request had a maximum of twenty turns.

### Web interface and output validation

Researchers interact with the framework through a web interface (Figure 1C): a request is typed in the query box and started with Run (1); the Figures tab (2) lists and rebuilds the study figures; the Agent tab (3) shows the live tool graph, in which each specialist agent and its tools are placed on a rotatable 3D brain and light up while they run; and the language model serving the session is shown at (4). The interface runs a single language model with access to these tools, separately from the search loop described above.

Pydantic schemas defined the required output format, and agents returned a fixed JSON structure. If validation failed, the system sent one repair request with the error message. A second failure stopped the run. The provenance log stored token use, prompts, model replies, tool calls, and validation results. This made it possible to measure cost and output failures. All reported numbers came from the deterministic tool layer. Before an agent output was used in the next step, the system replaced its numerical fields with the values in the tool record. This prevented an incorrect number produced or copied by the language model from entering the workflow.

### Regional and connection level analysis

We also examined where the classification signal was located. Connectivity strength was the mean absolute Fisher-z correlation between one region and all other regions in the same atlas. For each region, we tested its association with diagnosis while controlling for age, sex, motion, and center. We applied Benjamini–Hochberg correction within each parcellation. Missing coverage was handled separately for each region, so participants with incomplete coverage were not removed from the whole analysis. We used the same six parcellations as the prediction model. The anatomical analysis therefore describes the feature space used by the classifier (Figure 5).

### Statistical analysis

Pooled AUC confidence intervals were obtained with the DeLong method and with a cluster bootstrap that resampled acquisition centers (2,000 resamples); the significance of the pooled AUC was assessed with 10,000 label permutations. Confidence intervals for the six sealed test centers used the Hanley–McNeil method. All models were compared on identical out of fold predictions: the 24 ranked models were compared with the selected design using Wilcoxon tests with Holm correction (Figure 3B), and differences in AUC between ensembles were estimated with a paired cluster bootstrap over centers (2,000 resamples) with one sided P values for an improvement. Mean framewise displacement was compared between groups with the Mann–Whitney test, and associations were assessed with Spearman correlation. Motion matched pairs were formed within centers with a caliper of 0.2 s.d. of mean framewise displacement. Regional and connection level effects were estimated with regression models adjusted for age, sex, mean framewise displacement and center, with Benjamini–Hochberg correction within each parcellation.

## Resource availability Lead Contant

Requests for further information and resources should be directed to and will be fulfilled by the lead contact.

## Material Availability

This study did not generate new unique reagents.

## Data and Code Availability

ABIDE I data and C-PAC derivatives are publicly available from the Preprocessed Connectomes Project (http://preprocessed-connectomes-project.org/abide/). Out-of-fold predictions and the public subject identifiers used in this study are included in the code repository.

Code to reproduce all analyses is available on GitHub (https://github.com/karimrahimian/ana) and Figshare (https://doi.org/10.6084/m9.figshare.34866567).

## Acknowledgements

This research received no specific funding.

K.R, M.M contributed to data collection, K.R, R.B contributed to study design, K.R, MM design workflow and code, data analysis. K.R, S.B , data visualization. K.R and M.M wrote the manuscript, K.R, R.B monitored the accuracy of data. S.B designed graphical contents. L.H. Y.D have edited and provided useful comments. Y.D supervised the work.

## Declaration of Interests

### Declaration of generative AI and AI-assisted technologies in the manuscript preparation

The authors declare no conflicts of interest.

OpenAI ChatGPT assisted only with grammar, syntax, clarity and readability. The writing tools were not used to generate data, perform statistical analyses, interpret results or draw scientific conclusions. The authors reviewed and edited all resulting text and accept responsibility for the manuscript.

## Supplemental Information

S1-S5 are provided in the supplemental file: Table S1, ABIDE I sample by acquisition-site label; Table S2, contribution of each parcellation to the ComBat ensemble; Table S3, center-specific performance of the agent wavelet search; Table S4, rs-fMRI feature taxonomy used by the Literature Agent; Table S5, complete search log of the pipeline evaluations.

